# Multiplexed electrokinetic sensor for detection and therapy monitoring of extracellular vesicles from liquid biopsies of non-small-cell lung cancer patients

**DOI:** 10.1101/2021.04.08.438994

**Authors:** Sara Cavallaro, Petra Hååg, Siddharth S. Sahu, Lorenca Berisha, Vitaliy O. Kaminsky, Simon Ekman, Rolf Lewensohn, Jan Linnros, Kristina Viktorsson, Apurba Dev

## Abstract

Liquid biopsies based on extracellular vesicle (EV) protein profiles represent a promising tool for treatment monitoring of tumors, including non-small-cell lung cancers (NSCLC). In this study, we present the development of an electrokinetic sensor for multiplexed surface protein profiling of EVs and analysis of clinical samples. The method detects the difference in the streaming current obtained as a result of EV binding to the inner surface of a functionalized microcapillary, thereby estimating the expression level of a surface marker. Using multiple microchannels functionalized with different antibodies in a parallel fluidic connection, we first demonstrate the capacity for simultaneous detection of multiple surface markers in small EVs (sEVs) from NSCLC cells. To investigate the prospects of liquid biopsies based on EVs, we then apply the method to profile sEVs isolated from the pleural effusion (PE) fluids of three NSCLC adenocarcinoma patients with different genomic alterations (ALK-fusion, KRAS and EGFR) and applied treatments (chemotherapy, EGFR or ALK tyrosine kinase inhibitors). These vesicles were targeted against CD9 tetraspanin, as well as EGFR and PD-L1, two markers of interest in NSCLC. The electrokinetic signals showed detection of these markers on sEVs yet highlighting distinct interpatient differences, e.g., increased EGFR levels in sEVs from a patient with EGFR mutation as compared to an ALK-mutant one. The sensors also detected differences in PD-L1 expressions, in line with those measured by complementary methods. The analysis of sEVs from a patient prior and post crizotinib treatment also revealed a significant increase in the expression of some markers, e.g. EGFR and PD-L1. The obtained results hold promise for the application of the method for tumor treatment monitoring based on sEVs from liquid biopsies.

## 1. Introduction

Despite decades of research and development, cancer remains a major challenge for the global healthcare, killing millions of people every year (Sung et al., 2021). While the understanding of tumour biology as well as the methods of clinical interventions have seen major improvements, particularly regarding targeted therapy approaches, the progress on diagnostic technologies has been relatively slow and inefficient. This has also been due to the reliance on tissue biopsy as a source of tumour material (Vaidyanathan et al., 2019). For a long time, tumor tissue biopsies have been the gold standard of cancer diagnostics and subtyping, yet they have limitations in treatment monitoring as they only provide a restricted information of the patient tumor burden, in addition to being invasive and not easily accessible in several cases (Vaidyanathan et al., 2019). Therefore, there has been an exponentially increasing interest in non-invasive sampling of body fluids, known as liquid biopsy (De Mattos-Arruda and Siravegna, 2021; Rolfo et al., 2018; Santarpia et al., 2018). This is because such sampling approaches offer a less invasive way for treatment monitoring. Furthermore, liquid biopsies also give access to a plethora of ultrasensitive biosensing technologies, which have improved diagnostic approaches to many pathological conditions, including cancer (Vaidyanathan et al., 2019).

The prospect of EVs as a source of tumor biomarkers in liquid biopsies has been highlighted in a large number of research articles (Li et al., 2021; Santarpia et al., 2018; Xu et al., 2018). Lung cancer (LC), one of the tumor malignancies which causes most cancer related deaths (Sung et al., 2021) and where re-biopsy is challenging, has obviously been a subject of interest for the EV-based diagnosis (Hoshino et al., 2020; Pasini and Ulivi, 2020; Sandfeld-Paulsen et al., 2016; Wan et al., 2018; Zhong et al., 2020). In particular, non-small-cell lung cancer (NSCLC), the major subtype of LC accounting for ∼80% of all LC (Planchard et al., 2018) and which has a 5 year survival rate of only ∼ 20% (according to Cancer.org), has drawn particular interest, both from the perspective of early stage detection as well as monitoring of therapy (Pasini and Ulivi, 2020). Diagnostic biomarkers for NSCLC have been identified in both EV-protein profiles (An et al., 2019) and RNA content (Tao et al., 2020), thereby opening scope for developing biosensing methodologies targeting either of the two biomarker types. While different biosensing techniques, e.g., lateral flow aptamer assay (Yu et al., 2020), surface plasmon resonance imaging (SPRi) array (Fan et al., 2020) and streaming current based sensor (Cavallaro et al., 2019), have recently demonstrated the capacity of surface protein profiling of NSCLC cell derived EVs, the scope for clinical sample analysis has been mainly performed by traditional proteomic-analyses and sequencing-based methods (Hoshino et al., 2020; Hydbring et al., 2018; Krug et al., 2018; Sandfeld-Paulsen et al., 2016). A major challenge in analyzing EVs isolated from liquid biopsies of patients is the heterogeneity of the vesicles and the complex matrix of plasma, which also contains proteins that might get co-isolated with EVs (Ferguson and Weissleder, 2020). This means that, for such clinical applications, a biosensing method needs to have the capacity to measure multiple markers with a high specificity and reproducibility.

During the past years, the detection method employing the electrokinetic principles of streaming current/potential has been applied by various research groups to detect a variety of biomarkers (Dev et al., 2016; Koch et al., 1999; Martins et al., 2011). The method allows label-free detection of target molecules/bioparticles within a microfluidic channel by exploiting hydrodynamic and electrostatic interactions at the solid-liquid interface (Adamczyk et al., 2010). Moreover, the technique has a simple design and good sensitivity. In a previous study, we demonstrated the capability of this method for detection and profiling of EVs from cell lines (Cavallaro et al., 2019). In particular, the technique was able to discriminate changes in epidermal growth factor receptor (EGFR) expression in EVs isolated from a NSCLC cell line, with a sensitivity of 10% of the parental cell expression (Cavallaro et al., 2019). Since EGFR is often targeted in NSCLC using multiple tyrosine kinase inhibitors (TKIs) against it (Guo et al., 2020; König et al., 2021; Suda et al., 2017), the method holds a substantial promise for further validation in clinical samples. However, for such analysis, the development of a multiplexed assay is essential to simultaneously assess the changes in multiple EV surface markers among different patients and/or treatment stages. To the best of our knowledge, such capacity has not been demonstrated using the sensing method.

In this article, we first validated the possibility to use a multiplexed electrokinetic platform based on streaming current (Is) for the detection of EVs. For the purpose, we used vesicles isolated from the cell culture media of NSCLC cells and characterized the platform for the simultaneous measurements of four microcapillary sensors. Following validation, we applied the multiplexed platform for detection and profiling of EVs isolated from malignant pleural effusion (PE) fluids of three NSCLC patients with different genomic mutations and during different stages of their treatment courses. PE refers to the accumulation of fluid in the space between the lungs and the chest walls in both malignant and non-malignant conditions (Baburaj et al., 2020; Stiller et al., 2021). With respect to advanced LC patients, PE fluid accumulation is seen in about 10-15% of all cases with a predomination of adenocarcinomas due to their growth pattern in the lung (Baburaj et al., 2020). As malignant PE fluid is enriched in tumor cells, might also contain tumor cell derived EVs and can be collected in a minimally invasive way, it represents a suitable source for liquid biopsies based on EVs. For the analysis, we targeted CD9 tetraspanin and two biomarkers of interest in NSCLC, namely EGFR and PD-L1, expressed on the EV surfaces. The results demonstrated successful detection of EVs from PE samples, highlighting interpatient differences in the expression of the analyzed markers. Finally, aiming to examine the possibility to use the multiplexed electrokinetic platform for treatment monitoring, we analyzed the EVs from an ALK-positive NSCLC patient at two time points during the treatment course. Particularly, we compared the expression levels of CD9, EGFR, PD-L1 and HER2 on EVs from PEs collected prior and post Crizotinib treatment. The data suggested increased levels of some of the analyzed markers, e.g., EGFR and PD-L1, while showing stable expressions of other ones, e.g., CD9.

## 2. Materials and Methods

### 2.1. Reagents

High purity deionized water (DIW) with a resistivity of 18 MΩ·cm was used throughout all the experiments. Phosphate-buffered saline (PBS, P4417) in tablets was purchased from Sigma-Aldrich. Cetuximab antibody (Erbitux infusion solution, 5 mg/mL) was used to target EGFR and was purchased from Merck Serono. Anti-CD9 (MEM-61), anti-CD63 (mab5048), anti-HER2 (HRB2/258), anti-IGF1R (1H7), anti-PD-L1 (MAB1561) and isotype control (MAB002) antibodies were purchased from Bio-Techne. If not stated otherwise, all the other chemicals were purchased from Sigma-Aldrich Sweden AB.

### 2.2. sEV collection and isolation

The EVs investigated in this study were collected from different sources and isolated in order to obtain small EVs (sEVs, ∼30-300 nm). The sEVs used to characterize the multiplexed platform were collected from the cell culture media of a NSCLC cell line H1975 (ATCC^®^ CRL-5908™, LGC Standards, Teddington Middlesex, United Kingdom) with mutations in *EGFR (exon 20, T790M* and *exon 21, L858R)*(Pao et al., 2005). The sEVs from NSCLC patients were isolated from malignant PE fluids and named as PE002, PE009 and PE011. The patient tumours contained genomic alterations in *EML4-ALK* variant 3 (a/b) (PE002), *KRAS* exon 2, codon 12/13 (PE009), and *EGFR* exon 21, L858R (PE011), respectively. These samples were collected at Karolinska University Hospital, Stockholm, Sweden. The study was approved by the Ethics Review Authority in Sweden (https://etikprovningsmyndigheten.se), region Stockholm (EPN No. Dnr. 2016/2585-32/1, approval date 8^th^ of March 2017), with patient informed consent to study participation. The study was also approved with respect to tumor material into biobank and transfer for analyses at Uppsala University by Material Transfer Agreement. Both the NSCLC cell line and malignant PE fluid EV samples were isolated by size exclusion chromatography (SEC) using qEVoriginal Columns (Izon Science, Oxford, United Kingdom). In particular, the isolation of the sEVs from the cell culture media of NSCLC cells followed the protocol reported in our earlier studies (Cavallaro et al., 2019, 2021). For the PE fluid sEVs, we followed the procedure described in another earlier work (Stiller et al., 2021).

### 2.3. sEV characterization

The sEVs from cell culture media of NSCLC cells were characterized by nanoparticle tracking analysis (NTA) for their size and particle count estimations. As presented in Figure S1, the vesicles showed the expected size profile. In our earlier study, the same sEVs were further characterized by scanning electron microscopy (SEM), showing the expected vesicle sizes and morphologies (Cavallaro et al., 2021). Western Blot (WB) analysis was performed to verify the presence of sEV markers, e.g., CD9, EGFR and TSG101, and the absence of cellular contamination by calnexin, as previously described (Cavallaro et al., 2019). The expression of PD-L1, HER2 and IGF-1R in sEVs was examined by proximity extension assay (PEA) due to the higher sensitivity of the method as compared to WB. The analysis was performed on the multiplex Immune Oncology and Oncology II® panel (Olink Proteomics AB, Uppsala, Sweden) carried out on the Clinical Biomarker Facility, Science for Life Laboratory, Uppsala University, Uppsala, Sweden. The PEA assay consists of antibody pairs against protein biomarkers linked to different tumor or immune signaling processes (https://www.olink.com/products) and also harbor internal positive and negative controls of the assay. For analyses, sEVs were lysed in 5x RIPA buffer to a final concentration of 1x RIPA (50mM Tris-HCl pH 7.4, 150mM NaCl, 1% Triton-X100, 1mM EDTA, 0.1% SDS) from which 1µL was applied in the assay. Data processing was carried out on the Olink Wizard for GenEx software with the normalized protein expression (NPX) values used in subsequent analyses. The fold levels relative to each marker’s limit of detection (LOD) value are given for the samples. The protein characterization results by WB and PEA are presented in Figure S1. The size profiles and concentrations of the EVs from the human PE fluids were also characterized by NTA. For their biomarker characterization, WB (for EGFR, CD9, no calnexin) and proximity extension assay (PEA, for PD-L1) were used. The results of the analyses are reported in an earlier study ((Stiller et al., 2021), NTA profiles and WB) and in Supplementary Information (Figure S2). The details on NTA and WB analyses are provided in our earlier studies (Cavallaro et al., 2021; Stiller et al., 2021).

### 2.4. Multiplexed electrokinetic platform

Figure 1 shows a schematic of the measurement process, from the sEV isolation and characterization to the electrokinetic detection and profiling. Following isolation and characterization, electrokinetic experiments were performed in a platform consisting of a commercial pumping system (Elveflow, OB1), a flow sensor (Elveflow, MFS3), a fluidic system including the sensors, and a measurement unit (Keithley source meter and PC). The fluidic system comprised microfluidic tubing to transport the buffer/analytes from the sample vials to the sensors, a multiport connector for multiple sensor connections, silica microcapillary sensors and hollow Pt electrodes at the inlet and outlet of the microcapillaries. The electrokinetic measurements were performed in silica microcapillaries having inner diameters of 25 μm and lengths of ∼4.7 cm. A common Pt electrode was used for all sensors and connected at their inlet (before multiport connector), while separate Pt electrodes were connected to the different outlets. For each microcapillary, the current between the Pt electrode at the inlet and the one at the outlet was measured. A schematic of the platform is presented in Figure 1.

**Figure 1.**
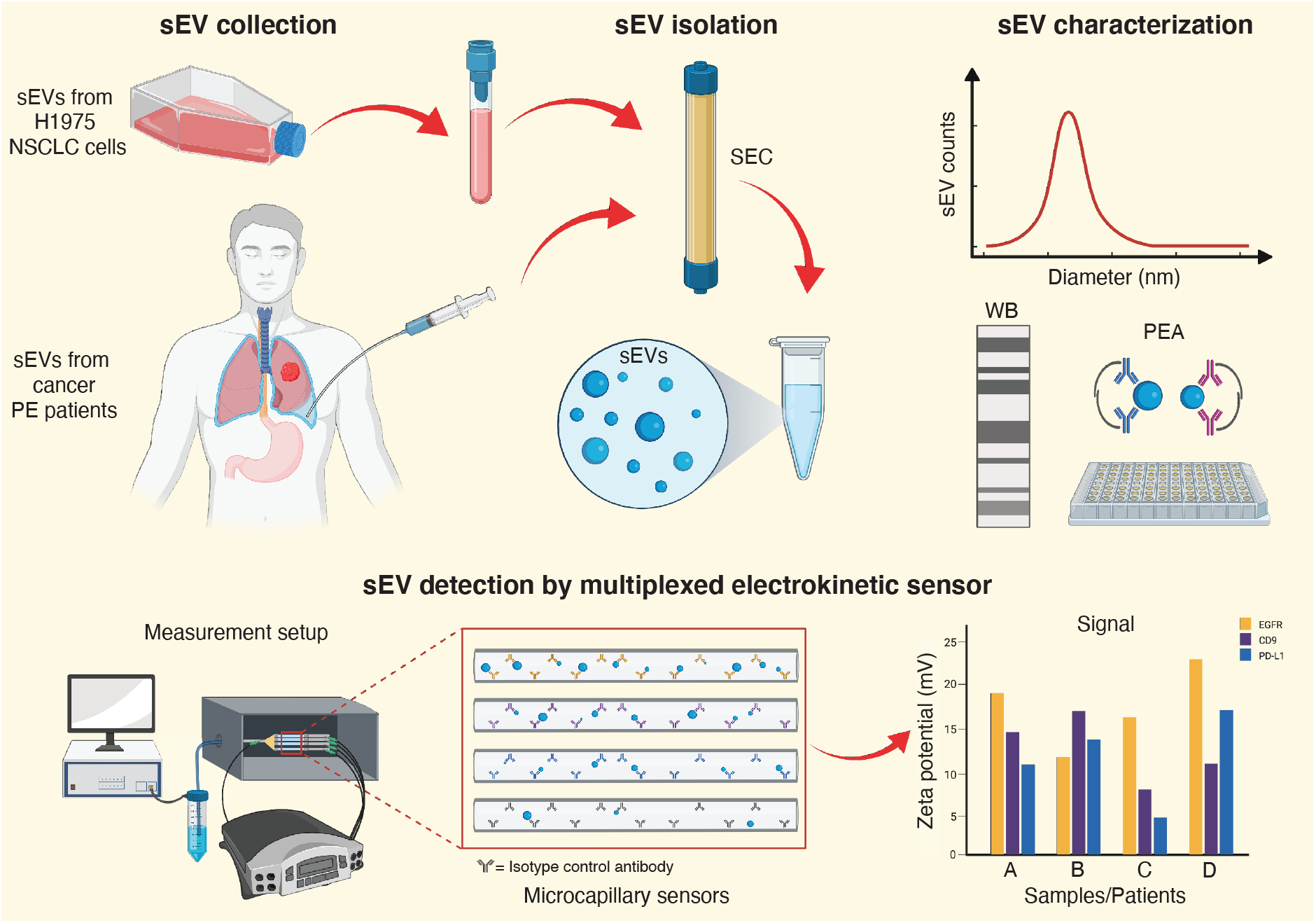
Schematic of the steps performed during the study. EVs from either cell culture media of NSCLC cells or malignant PE fluids of NSCLC patients were first isolated by SEC. The isolated vesicles were then analyzed by NTA for their size and concentration estimations, and by WB and PEA for their protein content characterization. The EV detection and profiling was performed by an electrokinetic platform consisting of 3 or 4 microcapillary sensors functionalized with different capture probes and connected simultaneously. The signals for the different targeted proteins were reported as changes in the zeta potential between second and initial baselines and compared among different samples. This figure was realized using BioRender.com.

The microcapillary sensors were functionalized following our previously reported protocol (Cavallaro et al., 2019). Briefly, after cleaning, (3-aminopropyl)triethoxysilane (APTES) activation and glutaraldehyde (GA), the antibodies targeting the different EV surface proteins were immobilized to the inner microcapillary surfaces for 2 h, at a concentration of 50 μg/mL in 1x PBS. Tris-Ethanolamine (30 min) and casein (120 min) were then used to deactivate the surfaces, thus reducing non-specific binding (NSB) of the vesicles (Cavallaro et al., 2019). The control capillaries were functionalized by following all the same steps, but immobilizing isotype control antibodies at the same concentration (50 μg/mL in 1x PBS), and were used to test the EV NSB. Prior to measurements, all microfluidic tubing were also treated with casein for 120 min to minimize NSB of the vesicles.

For detection, we analyzed the Is induced by a pressure driven flow of a 0.1x PBS buffer and measured the current changes upon EV binding to the functionalized microcapillary surfaces, as reported in our previous study (Cavallaro et al., 2019). In brief, for each channel we measured the Is for a continuous periodic (30 s pulse duration) rectangular pulse of 1.5 bar pulse height (1.5 bar and 3.0 bar applied pressures), which was used to drive the PBS solution through the microcapillaries. The current difference (ΔIs) for the applied pressures was recorded by the source meter and converted in apparent zeta potential (ζ*), according to the equation:

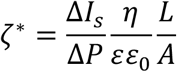

Where ΔIs/ΔP is the change in Is with pressure, η the viscosity, εε_0_ the permittivity, and L/A the length and cross-section of the capillaries, respectively. The collected data were analyzed by a custom-designed MATLAB software. All experiments were performed at room temperature with 0.1x PBS at pH 7.4, following an end-point detection approach, as described earlier (Cavallaro et al., 2019). Specifically, two baselines were recorded, one prior to and one post EV incubation in the microcapillaries, and the signals were calculated as the difference in the ζ* between these two baselines (Δζ*).

## 3. Results

### 3.1 Multiplexed platform validation with EVs from NSCLC H1975 cells

To setup and characterize the multiplexed electrokinetic platform, we first tested the effects of multichannel connection on the measurement parameters. In particular, we compared the noise in current measurements and the changes in the baseline values upon connection of 1 to 4 simultaneous microcapillary sensors. We selected a maximum of 4 parallel channels for the proof of principle study, as this could be easily setup by using commercially available manifold connectors. Figure 2A shows the noise in the ΔI_s_, calculated as the standard deviation of a blank, for microcapillaries in different channel configurations. For easier comparison with the following data reported in the study, we also included the corresponding noise levels in ζ* (Figure 2A, inset bar plots). For the noise analysis, we considered microcapillaries functionalized with the different capture probes reported in Materials and Methods. As visible in Figure 2A, the current noise in the two-channel configuration remained the same as the single-channel measurement (∼0.1 pA), while showing a 1.5-and a 2-fold increase in presence of 3 and 4 simultaneous channels, respectively. To test the presence of cross-interference between the different channels, we further measured the changes of the initial baseline values (ζ* prior to EV incubation) upon connection of 2, 3 and 4 simultaneous capillaries, with respect to their corresponding single-channel baselines. Figure 2B shows these results for a CD9-antibody functionalized sensor and a control one. As a reference, we set the corresponding initial single-channel baselines as ζ*=0 mV and reported the changes in absolute ζ* values.

**Figure 2.**
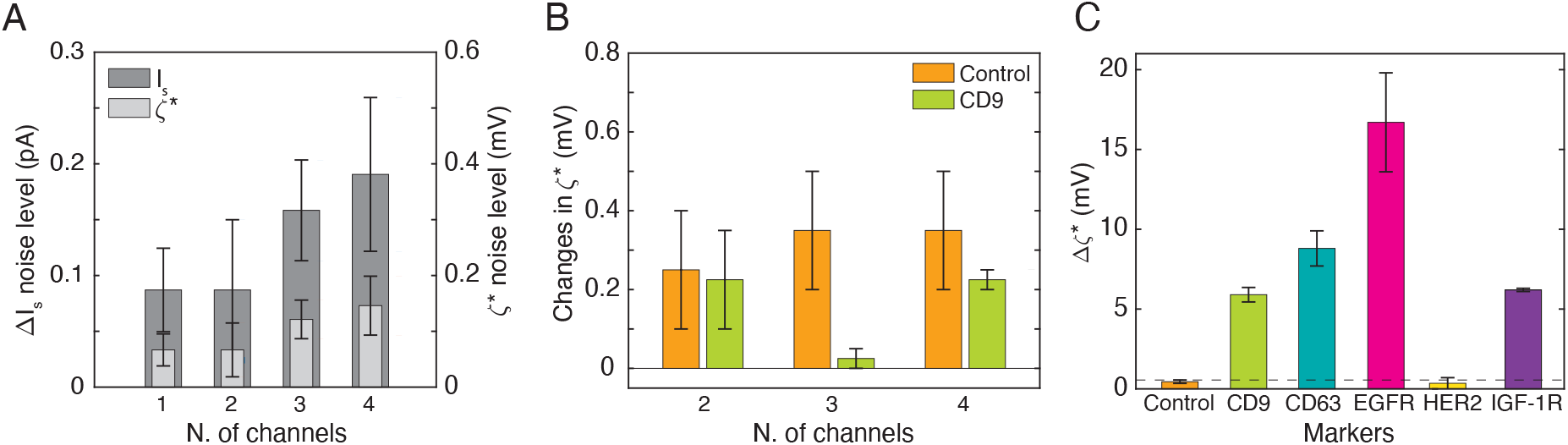
Multiplexed electrokinetic platform characterization and validation with sEVs from conditioned cell culture media of NSCLC H1975 cells. (A) Noise levels in the ΔI_s_ (left y-axis) and corresponding ζ* (right y-axis) for single- to four-channel configuration measurements, calculated for 5 different microcapillaries. (B) ζ* changes in the initial baselines (in absolute values) when measuring two to four simultaneous channels as compared to the corresponding single-channel measurements. An EGFR-functionalized and a second control capillary were connected as 3^rd^ and 4^th^ channels (not shown in the plot). Measurements performed in duplicate. (C) Sensor responses for the different capture probes CD9, CD63, EGFR, HER2 and IGF-1R measured by the multiplexed platform. sEVs incubated in the microcapillary sensors at a concentration of 3.5 × 10^9^ particles/mL. Measurements performed in duplicate. The dashed line depicts the MDS of the platform in its four-channel configuration measurement (∼0.5 mV).

As presented, both sensors showed negligible baseline variations, in the order of the minimum detectable signals (MDSs, defined as three-times the noise levels, as defined above) of the corresponding numbers of channels. This suggested the absence of cross-interference, indicating that each capillary sensor could be measured independently of the others. However, we noticed a slightly unequal distribution of the buffer among the different microcapillaries when connecting 4 channels simultaneously (∼13% variation in net flow/min). The flow was instead distributed more uniformly (< 7% variation) when connecting 2 or 3 channels in parallel.

Following initial characterization, we tested our four-channel multiplexed platform with sEVs isolated from conditioned cell culture media of a NSCLC cell line H1975. For the analysis, we targeted the ubiquitous EV tetraspanins CD9 and CD63, and some biomarkers of interest in NSCLC therapy, e.g., EGFR, HER2 and IGF-1R. These tumor cell surface receptors are known to be oncogenic drivers of NSCLC, with EGFR also being a precision medicine target and HER2 and IGF-1R both reported to be bypass drivers in EGFR-TKI refractoriness (König et al., 2021; Lai-Kwon et al., 2021). As presented in Figure 2C, all the functionalized sensors showed a positive response as compared to control capillaries (with isotype control antibodies), except for HER2. We could not detect this protein using the selected antibody and sEV concentration (3.5 × 10^9^ particles/mL). For comparison, we also analyzed the same surface markers using WB and PEA. The results are reported in Figure S1 and confirmed the expression of these proteins on sEVs from NSCLC H1975 cells.

### 3.2. Detection of sEVs from malignant PE fluids of NSCLC patients

Following characterization and validation with sEVs from NSCLC cells, we tested whether our multiplexed electrokinetic platform could detect EVs from patient samples and discriminate in their biomarker expression levels. Figure 3A summarizes details on the tumor types, stages at diagnosis and treatment courses at sample collection for the analyzed patients.

**Figure 3.**
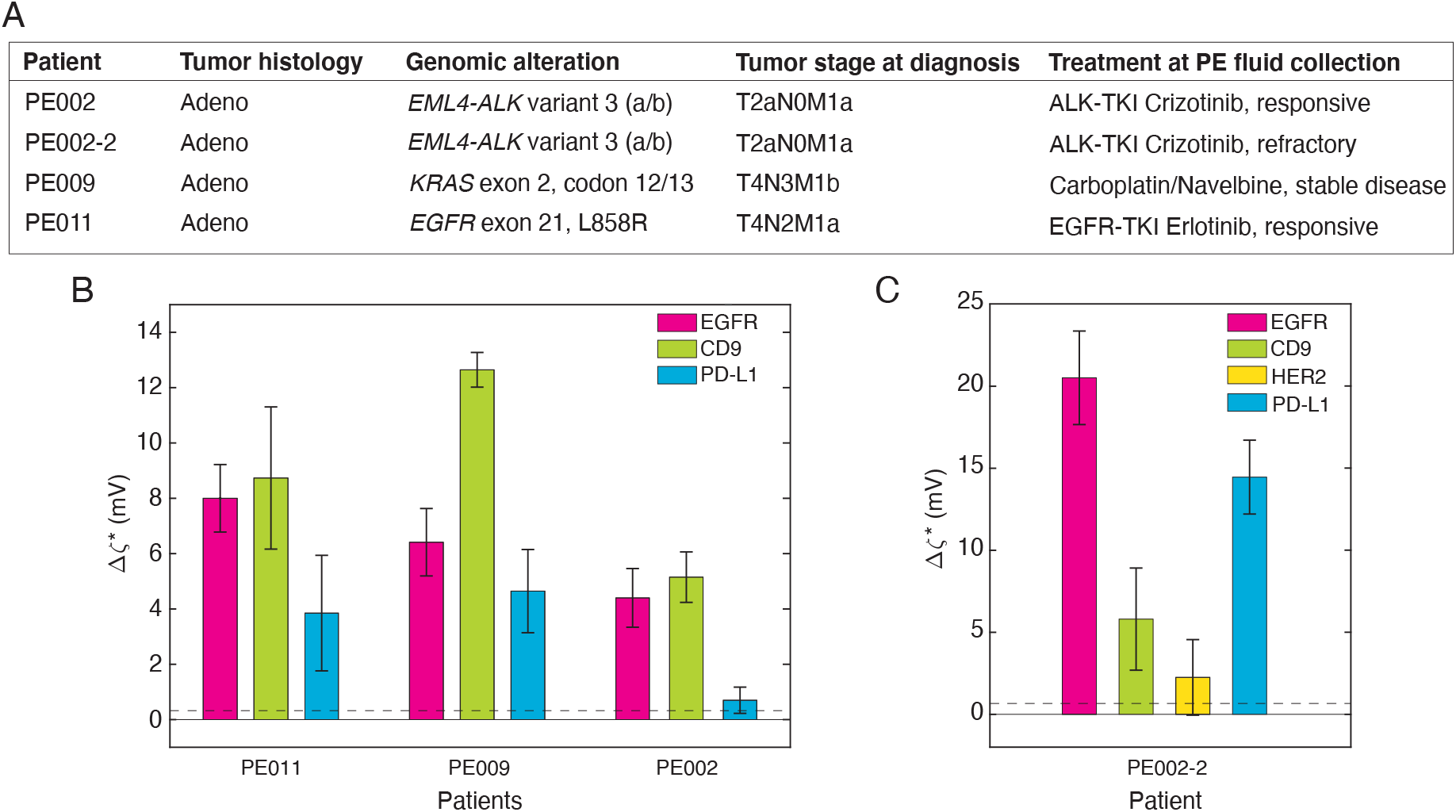
Multiplexed electrokinetic platform application to sEVs from malignant PE fluids of NSCLC patients. (A) Table summarizing the molecular and clinical data of the malignant PE fluids, including tumor histology, genomic alteration, tumor stage at diagnosis according to TNM 8th edition, treatment at time of PE collection and clinical response of the patients. (B) Bar plot showing the sensor responses when targeting the sEVs from PE002, PE009 and PE011 against CD9, EGFR and PD-L1. Measurements performed in triplicate, incubating the sEVs at a concentration of 3.5 × 10^9^ particles/mL. Controls (capillaries with isotype control antibodies) subtracted from each bar plot and sample. (C) Bar plots showing the sensor responses for the sEVs from PE002-2 sample targeted against the same markers as in (B). HER2 also analyzed. Measurements performed in duplicate. The dashed lines depict the MDS of the platform in its three-channel configuration measurement (∼0.4 mV).

In details, we analyzed the sEVs isolated from the PE fluid samples of three malignant adenocarcinoma NSCLC patients with different genomic alterations, namely *EML4-ALK* variant 3 (a/b) (PE002), *KRAS* exon 2, codon 12/13 (PE009), and *EGFR* exon 21, L858R (PE011), respectively. PE002 was on ALK-TKI crizotinib when PE sampling was obtained and responded to this treatment line. PE009 was on standard chemotherapy at the time of sampling with stable disease. PE011 was given EGFR-TKI erlotinib with a clinical response. As compared to cell line derived EVs, samples isolated from human body fluids might also contain impurities/biomolecules that may interfere with the detection method, therefore causing higher NSB. For electrokinetic detection, we investigated the transmembrane proteins CD9, EGFR and PD-L1 and compared their levels in the sEVs isolated from the three PE samples. We included PD-L1 in the analysis as it has been demonstrated that the tumor expression of this marker may impede on the immune system to attack the tumor, thereby offering a way to monitor therapy (Abdel Karim and Kelly, 2019; D. H. Kim et al., 2019). It has also been suggested that PD-L1 is expressed in EVs (Wang et al., 2021) and that Tyrosine Kinase Inhibitors (TKIs) against mutated EGFR may influence PD-L1 expression as well as the immune landscape in NSCLC (Kang et al., 2021; Liu et al., 2021). Moreover, for this investigation, we used the platform in a three-channel configuration, in order to reduce the inhomogeneous flow distribution among the capillaries that was detected when connecting 4 channels simultaneously. For comparison, we analyzed the same sEV batches for CD9, EGFR and PD-L1 using WB and PEA. These results are presented in an earlier work (Stiller et al., 2021) and in Figure S2. The bar plots in Figure 3B show the sensor responses (upon control subtractions) for the three patients and surface proteins analyzed. Control measurements (capillaries with isotype control antibodies) showed lower responses (Figure S3) than the respective signals. However, in some cases, increased NSB were detected (∼5.5-fold for PE009 and ∼4.8-fold for PE011) as compared to the controls obtained with sEVs from cell lines (∼0.4 mV, Figure 2C). As presented in Figure 3B, the electrokinetic data revealed positive detection of both CD9 and EGFR for all patients, but with a differential expression among the samples. In particular, EGFR was found to be more abundant in PE011 and PE009 (∼8 mV and ∼6.5 mV) as compared to PE002 (∼4.4 mV). CD9 instead, seemed to be more abundant in PE009 (∼12.5 mV) as compared to PE011 (∼8.7 mV) and PE002 (∼5.2 mV). The same trends were confirmed by WB profiling, which showed a higher expression level of both these markers on sEVs from PE011 and PE009 as compared to PE002 (Figure S2 and (Stiller et al., 2021)). With respect to PD-L1, a higher electrokinetic response was evident in PE009 (∼4.6 mV) as compared to PE011 (∼3.8 mV), while the signal was just above the MDS for the sEVs from PE002 (∼0.7 mV, Figure 3B). The expression of this marker was also analyzed on the same sEV batches using PEA analytics (Figure S2). As seen, PEA results also revealed similar differences in PD-L1 expression levels among the analyzed samples.

### 3.3. Effect of treatment on sEV surface markers

Finally, to examine the potentials of the electrokinetic method for treatment monitoring, we further analyzed the sEVs from the PE002 patient collected after 1-year on ALK-TKI crizotinib treatment, when the patient was clinically refractory. We refer to this sample as PE002-2 (Figure 3A). Apart from compensatory mutations in the ALK kinase domain bypass signalling (Gristina et al., 2020), it is reported that ALK-TKI resistance also includes alterations in EGFR, IGF-1R and HER2 (Choi et al., 2017; Dagogo-Jack et al., 2017; Gristina et al., 2020; Minari et al., 2020). For this reason, we also included HER2 in addition to the previously studied markers CD9, EGFR and PD-L1 for profiling of sEVs from PE002-2. When HER2 was analyzed on the sEVs from the initial PE002 sample, the sensor did not show any response (data not shown). Figure 3C presents the sensor responses for the markers analyzed on sEVs from PE002-2 patient. With respect to CD9, similar average values (∼5 mV) were observed in sEVs from PE002-2 as compared to those detected at initial sample collection (PE002). On the contrary, EGFR and PD-L1 levels on sEVs significantly increased after crizotinib, showing a ∼4.5-fold and a ∼21-fold increase, respectively in PE002-2, as compared to their levels in sEVs from PE002. Finally, HER2 could be detected in PE002-2, despite showing a weak response (∼2 mV).

## 4. Discussion

A key benefit of liquid biopsies is the ability to follow cancer patients during their treatments, thereby guiding the clinician with effective therapy choices such as precision cancer medicine agents or immune checkpoint inhibitors (ICI). This is important also in NSCLC, where EGFR-TKI, ALK-TKI or PD-1/PD-L1 immune therapy have changed the treatment landscape, however responses of patients are often heterogenous (Abdel Karim and Kelly, 2019; Gristina et al., 2020; Lazzari et al., 2020). In clinical settings, circulating/cell-free DNA is partially used as a biomarker (BM) for evaluating treatment response when tumor biopsy is not feasible (Rolfo et al., 2018). However, DNA does not give information on alterations in tumor signalling taking place on mRNA or protein levels. In these cases, EVs from tumor cells *per se* as well as from immune cells within the tumor may offer a superior source of BMs. In this study, we present an electrokinetic biosensor that is capable of monitoring several sEV and tumor cell surface markers in sEVs isolated from PE fluids of NSCLC patients. However, to be used in clinical applications, a biosensor needs to have the capacity to measure multiple markers simultaneously from a low sample volume with a high specificity and reproducibility. The data on the platform characterization and with sEVs isolated from cell culture media of NSCLC cells (Figure 2) demonstrate that our electrokinetic sensor based on streaming current is suitable for multiplexed analysis of EV surface proteins. As shown in Figures 2A and 2B, the measurements parameters do not change significantly upon multiplexed connection of 4 microcapillary sensors. In particular, Figure 2A indicates that, despite a ∼2-fold increase when connecting more than 2 channels, the current noise still remains low (<0.3 pA) for three and four-channel configurations, as compared to the detected signals (>25-30 pA, data not shown). Such increased noise levels might come from the electrical measurement and/or from the fluidic part, due to flow splitting at the multiport connector and distribution among the different channels. Small variations in the ζ* baselines, as those reported in Figure 2B, might also be caused by small changes in the flow of one channel upon connection of the others. However, all such ζ* changes (Figure 2B) are in the order of the MDSs of the corresponding number of channels (Figure 2A) and therefore can be neglected. Overall, these data suggest that the measurement outcomes do not change whether a marker is measured alone or in combination with others in parallel. Here, we would also like to emphasize that we tested a maximum of 4 channels only due to the difficulties to connect more of them in parallel in the current platform. This is not a limitation of the electrokinetic method and therefore in principle such measurements can be performed with a higher number of channels. Figures 2C demonstrates the successful application of the multiplexed electrokinetic platform for analysis of sEV surface markers. The results also suggest that the signals can be enhanced by optimizing the capture probes, as also reported in our previous study (Sahu et al., 2020). In particular, CD9, EGFR and CD63 proteins showed higher responses as compared to our previous article (Cavallaro et al., 2019). A possible explanation is the fact that we used a different set of capture probes, which had a more positive charge as compared to the previous ones and the EVs, therefore leading to enhanced signals (see Figure S4 and Supplementary information). The low signals obtained for HER2 can be explained by a very low surface expression level of the marker in these sEVs (as compared to other sEVs and/or tumor receptors) and/or by a low affinity of the capture probe used. On the other end, the results with sEVs from NSCLC cells suggested the presence of slight differences in the flow profiles (Δflow for the applied pressures of 1.5 and 3 bar) among the different channels when connecting 4 capillaries simultaneously. This effect might be caused by differences in the fluidic microcapillary resistances and/or flaws in the multiport connector that distributes the flow among the channels. Therefore, for the present investigation, we decided to measure the EVs from the malignant PE fluids of patients with the platform in its three-channel configuration.

Furthermore, the measurements were performed with sEVs at a concentration of 3.5 × 10^9^ particles/mL and required ∼25 µL of sample per microcapillary sensor. This corresponded to a total sample volume of ∼100 µL for a four-channel configuration platform, and therefore a total of 3.5 × 10^8^ EVs. For the PE samples used in the present study, we isolated EVs from 0.3-5 ml of filtered PE fluid. The number of EVs per ml of PE fluid was ∼1 × 10^10^, meaning that only <30-40 µL of liquids is necessary to perform such a multiplexed measurement with the proposed electrokinetic platform.

The results on the clinical samples (Figure 3) demonstrate the possibility to use the platform for detection and treatment monitoring of sEVs collected from PE fluids of advanced NSCLC patients. A challenge arising when analyzing patient samples can be the presence of proteins and/or analytes that might get co-isolated with EVs and interact with the biosensor, thereby causing high NSB. However, our data suggest that, despite an increase as compared to cell line derived vesicles, the sensor responses for control measurements on EVs from patient liquid biopsies were still significantly lower than those detected for the different EV markers. Since, the PE samples were collected from patients with different genomic drivers of their tumors, e.g., EGFR, ALK and KRAS, whom also have been given different treatment regimens including EGFR- and ALK-TKIs (Figure 3A), we expected different expression levels of the analyzed markers and therefore different responses of the multiplexed electrokinetic sensors. The results confirmed the expectations. For example, the sEVs from PE011, which were obtained from a patient with EGFR-mutant NSCLC, showed a stronger EGFR signal as compared to the sEVs from PE002, which were instead collected from a patient with an ALK-driven tumor. We detected intermediate levels of EGFR on the sEVs from PE009, whose patient had instead a KRAS mutation. With respect to CD9, PE002 also showed the lowest levels among the analyzed PE fluids. The analyses of EGFR and CD9 by WB ((Stiller et al., 2021) and Figure S2) confirmed the same trends, demonstrating that our electrokinetic data is in line with the results obtained by standard techniques.

In our study we also analyzed PD-L1, an important ICI target in NSCLC, which also have been found and have a function in sEVs from NSCLC patient liquid biopsies (D. H. Kim et al., 2019). While the electrokinetic sensors detected a clear signal from PD-L1 on sEVs from PE011 and PE009, the vesicles from PE002 showed instead very weak levels. When PD-L1 expression was evaluated with PEA analytics, there was a clear difference in expression among PE002, PE009 and PE011 sEVs, confirming the sensor responses (Figure S2). These results suggest different PD-L1 levels on sEVs from malignant PE fluid samples that can also be captured by our electrokinetic sensors. Moreover, it has been shown that EGFR-TKI refractory NSCLC cells have a higher PD-L1 expression level (D. H. Kim et al., 2019) and such PD-L1 expressing sEVs were also reported to be functionally active. It was also recently demonstrated that PD-L1 expression in *EGFR*-mutated NSCLC is a bad prognostic factor (Kang et al., 2021). With respect to these results, it would hence in the future be interesting to analyze PD-L1 expression in sEVs by our sensor prior and post EGFR-TKI treatment of NSCLC patients.

Finally, the analysis of PE002-2 revealed that both EGFR and PD-L1 sensors gave a higher signal at the time point of ALK-TKI crizotinib refractoriness than early upon treatment induction (PE002), suggesting that crizotinib treatment might cause increased expressions of both these receptors in sEVs. HER2 also showed a higher response than those detected in PE002, yet the increase was less prominent (∼2-fold). Such increased EGFR or HER2 expression/functional dependency has been reported in literature for ALK-TKI refractory NSCLC cells and patients (Dagogo-Jack et al., 2017; Gristina et al., 2020; Minari et al., 2020). Albeit ICI treatment of EGFR- or ALK-driven NSCLC has been reported to be lower than in NSCLC in general (Gainor et al., 2016), it has been shown that PD-L1 expression is increased in ALK-TKI *in vitro* NSCLC models (S. J. Kim et al., 2019). The electrokinetic sensors also detected significantly increased levels of PD-L1 on the sEVs from the ALK-TKI refractory patient PE002-2. However, further analyses of sEVs from a larger cohort of patients, in general, and ALK-TKI treated patients, in particular, are needed to make general conclusions on possible cancer resistance mechanisms based on EV measurements.

## 5. Conclusions

In conclusion, in this study we first demonstrated the possibility to use the electrokinetic method to perform multiplexed detection and profiling of sEV surface proteins. The results on the platform characterization (Figure 2) indicated that the measurement parameters remain stable upon multiplexing, despite some unequal flow distributions when connecting 4 channels simultaneously. However, we believe that this limitation will be solved in the future, when bringing the platform on a chip, as the channel design and dimensions can be precisely optimized and controlled, in order to measure more than four EV surface proteins simultaneously. The results on the vesicles isolated from the PE fluids of different NSCLC patients (Figure 3) demonstrated the capability of the electrokinetic sensor based on streaming current to profile several tumor cell surface markers on EVs from clinical samples. Although we cannot state any clear conclusion on the relation between the analyzed proteins and the possible cancer resistance mechanics, due to the limited number of PE samples analyzed, the data indicated interpatient differences. Moreover, the electrokinetic results were shown to be in line with the results obtained by standard techniques, such as WB and PEA. This gives hope that in the future, following application to a larger sample cohort, the platform can be potentially used for monitoring the patient responses during the cancer treatment courses.

In the future, we plan to bring the platform on a chip, in order to reduce its size and further enhance its multiplexing capability. In this way, we will be able to scan a larger cohort of patients and markers in a faster way, thereby trying to find their relations with resistance/responsive mechanisms. We are also working on the functionalization strategy in order to enhance the charge difference between the EVs and the sensor surfaces, therefore increasing the signals and improving the LODs. Finally, we will explore the possibility to use the electrokinetic method for sandwich detection, to analyze multiple surface markers simultaneously in the same microcapillary and improve EV detection in complex samples, e.g., plasma.

## Author contributions

SC: conceptualization, data curation, formal analysis, investigation, validation, visualization, writing – original draft, writing – review and editing

PH: data curation, formal analysis, resources, writing – review and editing

SSS: data curation, formal analysis, software, writing – review and editing

LB: data curation

VK: data curation, writing – review and editing

SE: resources, investigation, project administration

RL: resources, funding acquisition, project administration

JL: funding acquisition, project administration, writing – review and editing

KV: resources, funding acquisition, writing – original draft, writing – review and editing

AD: method development and conceptualization, supervision and funding acquisition, project administration and writing – original draft, writing – review and editing

## Declaration of competing interest

The authors declare no competing interests.

## Acknowledgements

This study was supported by grants from the Erling Persson Family Foundation, Stockholm Cancer Society (#171123, #191293, #201202,) the Swedish Cancer Society (CAN 2015/401; CAN 2018/597), Stockholm County Council (#20160287, #20180404) and funds of Karolinska University Hospital FOUU (#75032.). For the tumor related part, the support and help from MSc Vasiliki Arapi, Dr Caroline Kamali, Dr Luigi De Petris, Dr Metka Novak and Dr Per Hydbring is acknowledged.

## Supplementary data

A supplementary file including characterization data and additional measurements in provided as a .pdf file.

## Notes

### Competing Interest Statement

The authors have declared no competing interest.

