## Supplementary file for "Multiplexed electrokinetic sensor for detection and therapy monitoring of extracellular vesicles from liquid biopsies of non-small-cell lung cancer patients"

**Figure S1**.

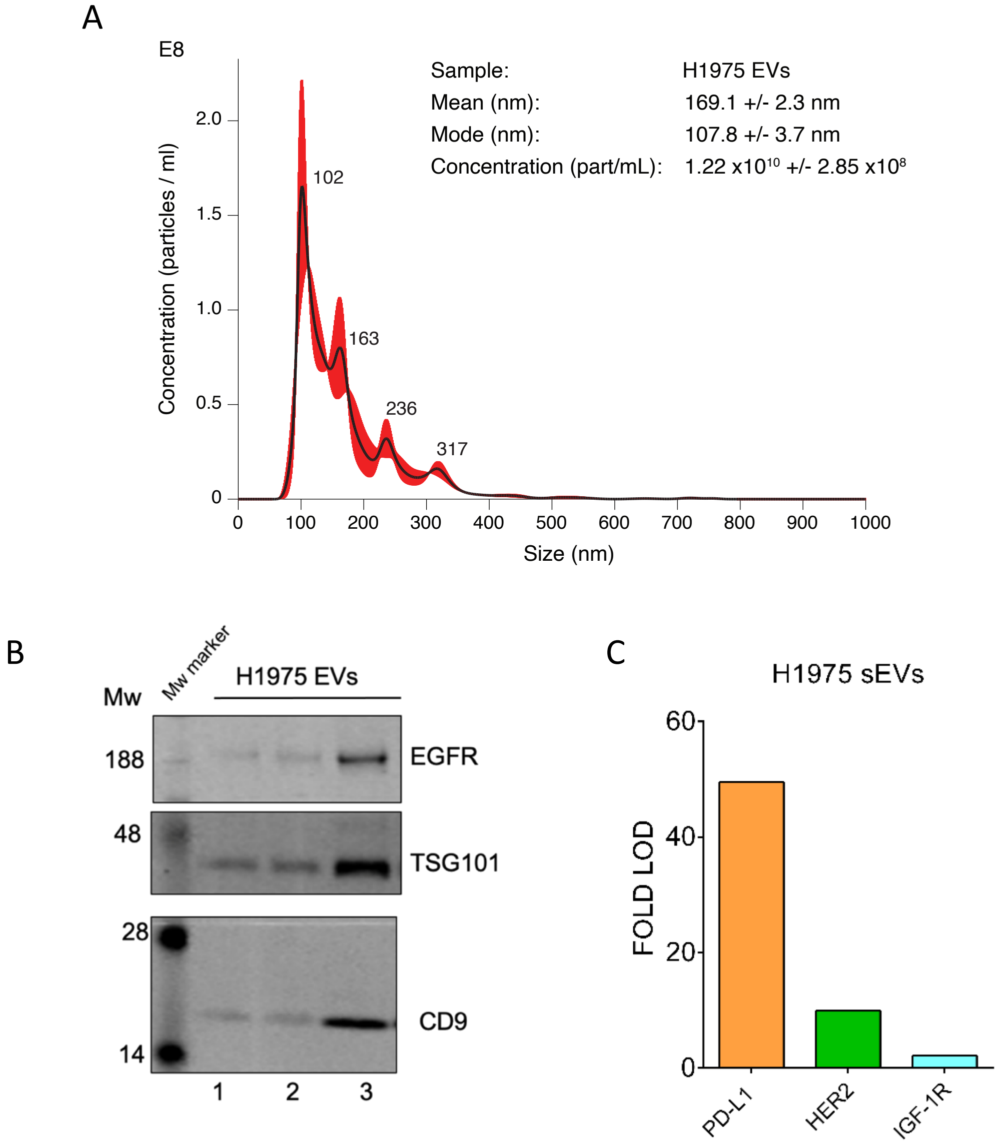


**Figure S1.** Characterization of sEVs from conditioned cell culture media of NSCLC H1975 cells. (A) NTA profile of EVs analyzed for mean and mode size, showing isolation of small EV sized vesicles (sEVs). Error bars indicate +/- 1 standard error of the mean. (B) Western blot analysis of sEVs from a replicate sample presented in (A) showing the presence of EV markers, e.g. CD9, TSG101, the tumor marker EGFR. The blot was also probed with the ER-protein calnexin which did not show any band (data not shown). The different columns correspond to the different amounts of EVs that were analyzed. In particular, line 1: 7,1x $10^7$ EVs, line 2: 1,4x $10^8$ EVs and line 3: 2,8 x$10^8$ EVs. (C) PEA analysis of HER2, IGF1R and PD-L1 levels on sEVs. The vesicles isolated from H1975 cell culture media were lysed in RIPA buffer and 1µl of sample, corresponding to 8.6 x $10^6$ EVs (as determined by NTA), was analyzed on PEA Oncology II and Immune Oncology platforms. The NPX values of the depicted markers were linearized and fold increase relative to limit of detection (LOD) values for each marker are shown.

**Figure S2**.

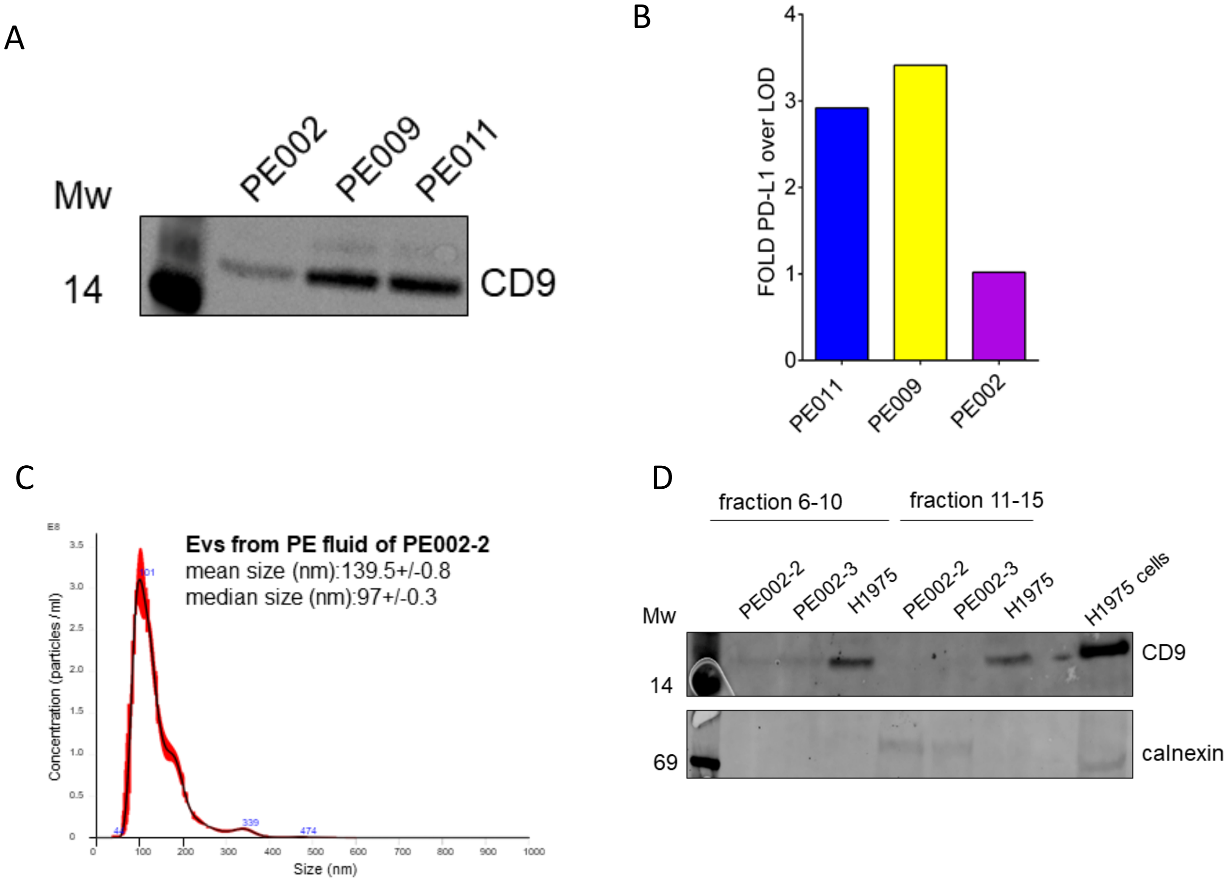


**Figure S2.** Characterization data for sEVs from PE samples. (A) WB analysis for sEVs from PE002, PE009 and PE011 for expression of CD9. The sEVs were lysed in RIPA buffert and equal amount of them (4.3 x $10^8$, as determined by NTA) was analyzed. (B) PEA analytics for PD-L1 analysis on sEVs from PE002, PE009 and PE011. Equal amount of sEVs was lysed in RIPA buffer and 1µl of sample, corresponding to 3.5 x $10^7$ (as determined by NTA) was analyzed on PEA Immune Oncology. The PD-L1 NPX values were linearized and fold increase relative to PD-L1 limit of detection is given. (C) NTA profile of sEVs from patient PE002-2 at 1-year following ALK-TKI crizotinib treatment. (D) Western blot analysis of EVs from PE002-

2 and another sample of the same patient PE002-3 taken after additional 4 months. The first 3 lines show samples from fraction 6-10 (corresponding to sEV sized EVs without protein contamination) of the qEV columns followed by fraction 11-15. As a control sEVs from condition media of H1975 cells and a cell extract of H1975 cells were applied. The membrane was probed by CD9 and the ER-protein calnexin.

The sEVs from PE002, PE009, and PE011 were the same as those utilized by Stiller et al. (Stiller et al., 2021). Therefore, the NTA profiles of these samples are reported in Figure S10 of the article by Stiller et al. (Stiller et al., 2021). WB analysis proving the presence of EGFR and the absence of cellular contamination (no calnexin) is also reported in the article by Stiller et al (Figure S11, (Stiller et al., 2021)).

**Figure S3**.

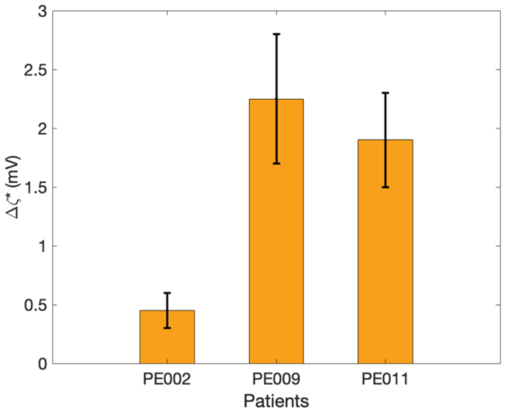


**Figure S3**. Control measurements performed in functionalized microcapillaries with isotype control antibodies for PE002, PE009 and PE011 samples.

**Figure S4**.

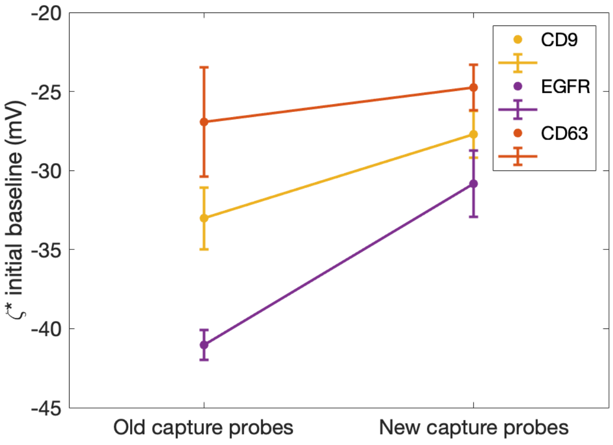

**Figure S4**. ζ* values of the initial baselines for different anti-EGFR, anti-CD9 and anti-CD63 antibodies as capture probes. Values calculated for 5 different microcapillaries. The capture probes utilized in this study, labelled as "new capture probes", have a less negative ζ* as compared to the probes used earlier in Cavallaro et al., 2019, labelled as "old capture probes".

Figure S4 shows the initial ζ* baselines for the new capture probes and the old ones utilized in our previous work (Cavallaro et al., 2019). In an earlier study, we demonstrated that when the surface has opposite charge as compared to the analyte, the electrokinetic signal is enhanced (Sahu et al., 2020). Since vesicles are negatively charged in PBS (ζ*~ - 40-45 mV) (Midekessa et al., 2020), we exploited this principle by using more positively charged CD9, EGFR and CD63 antibodies, compared to previous ones and EVs, that led to less negative initial ζ* baselines and therefore enhanced signals (Figure 2C) as compared to our previous work (Cavallaro et al., 2019).
